# Rim4 is a Thermal Sensor and Driver of Meiosis-specific Stress Granules

**DOI:** 10.1101/2024.01.09.574866

**Authors:** Rudian Zhang, Shunjin Li, Wenzhi Feng, Suhong Qian, Akshay John Chellappa, Fei Wang

## Abstract

Rim4 is a meiosis-specific RNA-binding protein (RBP) that sequesters mRNAs to suppress their translation. Previous work has defined the Rim4 C-terminal low-complexity domain (LCD) as sequences that form self-propagating amyloid-like aggregates. Here, we uncovered a dynamic and reversible form of Rim4 self-assembly primarily triggered by heat during meiosis, proportionally from 30°C to 42°C. The formed thermal Rim4 condensates in cell promptly stimulates stress granule (SG) assembly, recruiting SG-resident proteins, such as Pab1 and Pbp1, and strikingly, decreases the required temperature for meiotic SG formation (∼33°C) by ∼9°C as compared to mitosis (∼42°C). This sensitization of meiotic SG formation to heat effectively prevents meiosis progression and sporulation under harmful thermal turbulence. Meanwhile, the Rim4-positive meiotic SGs protect Rim4 and Rim4-sequestered mRNAs from autophagy to allow a rapid recovery from stalled meiosis upon the stress relief. Mechanistically, we found that the yeast 14-3-3 proteins (Bmh1 and Bmh2) and nucleic acids brake initiation of heat-induced Rim4 self-assembly, and Hsp104 facilitates the restoration of intracellular Rim4 distribution during the recovery.

## Main text

The subcellular membraneless compartments (MLCs) are driven by phase separation, also described as membraneless organelles (MLOs) featuring biomolecular condensates (∼0.2 to ∼2.0 µm in size) that typically consist of proteins and nucleic acids^1^. Unlike membrane-bound organelles, the biogenesis, regulation, and biological functions of MLOs are largely elusive. Remarkably, the nucleic acid-binding proteins, especially the RNA-binding proteins (RBP), are prone to be recruited to MLOs, thereby enabling the MLOs to function as platforms in the nucleus and the cytoplasm to regulate gene expression ^2–4^. Particularly, MLOs dynamics respond to the extracellular and intracellular cues, altering transcription and translation. Notably, the germline cells feature MLOs of specific functions; e.g., the nuage is an MLO regulating the mobile element ^5^; the P granule of > 50 proteins and various RNAs in *C. elegans* regulates germ cell-specific RNAs ^6^. However, little is known about MLOs during meiosis, the specialized cell cycle that leads to the generation of sexual cells.

Due to the scarcity of meiotic cells in higher eukaryotes, we initiated a project to identify MLOs of meiosis-specific RBPs that regulate meiosis, using *Saccharomyces cerevisiae* (budding yeast) as a model system. We found that Rim4, a meiosis-specific RBP known as forming amyloid-like aggregates essential for meiosis (depicted in Fig. 1A; 1B) ^7,8^, forms reversible stress granules (SGs). SGs are biomolecular condensates composed of RNAs and proteins when the cell is under stress ^1,9,10^. Particularly, the morphology of intracellular Rim4 changes from a mesh-like diffusion state to a spatially concentrated pattern under multiple stress conditions such as PH decrease in sporulation medium (PH 5.5) and oxidative stress (0.5% NaN_3_) but primarily by heat (37°C) (Fig. 1C). Strikingly, temperature increase between 30°C and 42°C stimulates Rim4 uneven distribution into discrete foci and proportionally (Fig. 1D-1F) leads to formation of cytosolic SGs marked by Pab1 and Pbp1 (Fig. 1G-H, S1A). Unlike the static amyloid-like aggregates previously reported, the Rim4 foci can rapidly disperse in ∼15 minutes once the stress was relieved to allow recovery (Fig. 1C). In addition, Rim4 that deletes one (ΔpolyN_1_ and ΔpolyN_2_) or two prion-like poly asparagine (ΔpolyN_12_) regions implicated in amyloid-like aggregation forms foci normally (Fig. S1B). Moreover, preformed Rim4 thermal foci partially disassemble in detergent (0.5% Triton X100)-ruptured cells, and fully disappear after 10 minutes of 2% SDS treatment at 23°C (Fig. S1C), indicating that these cytosolic Rim4 foci represent dynamic SG assembly instead of static amyloid-like aggregates previously reported ^7,11^. As a further support from Fluorescence *in situ* Hybridization (FISH), we found that mRNAs are recruited to cytosolic Rim4- and Pab1-positive foci under thermal stresses (Fig. 1I), a feature of cytosolic SGs.

**Figure 1.**
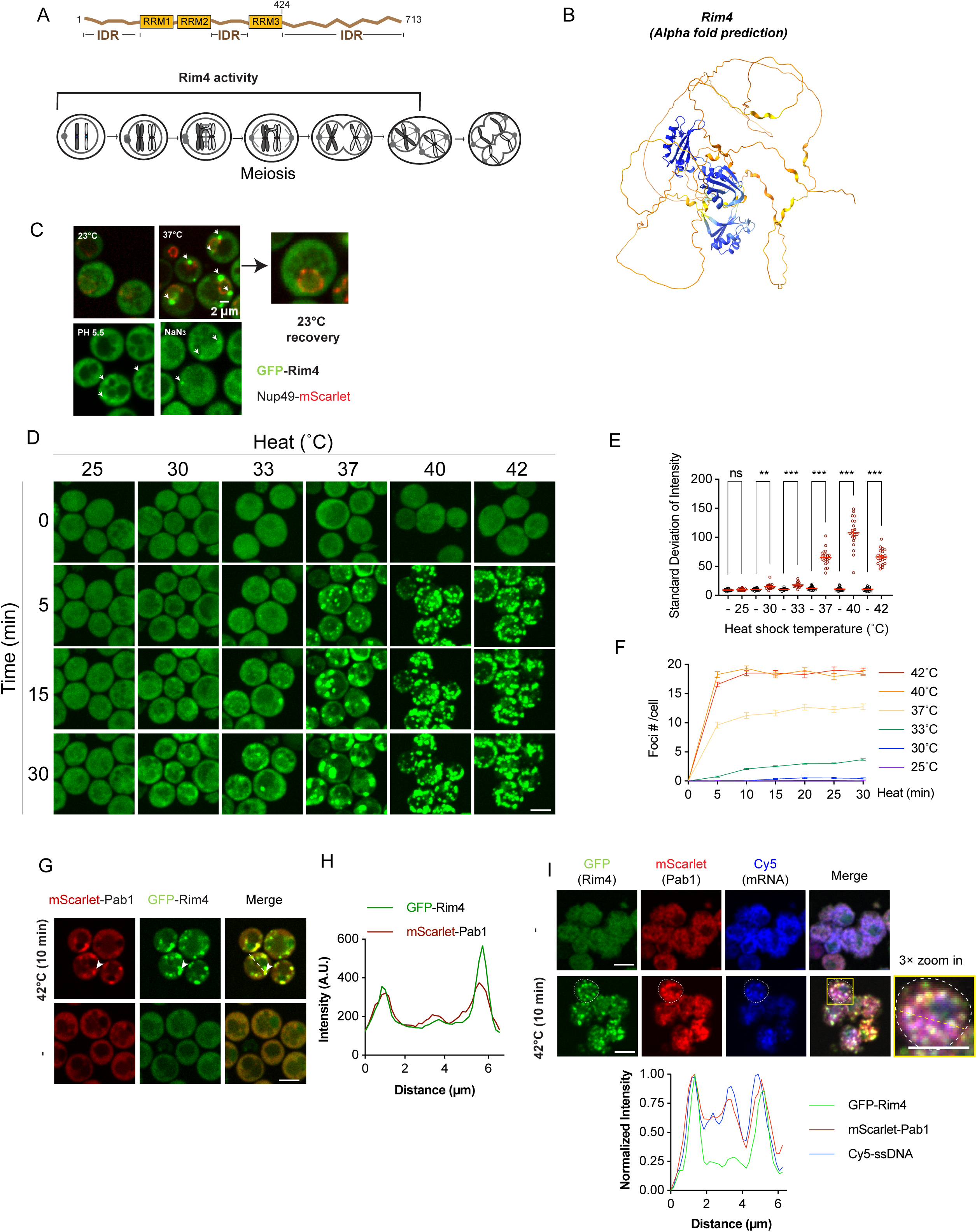
Rim4 is sensitive to heat and localizes to heat-induced stress granules (SGs) **(A)** Top: schematic depicts the domains of the Rim4 protein. RRM: RNA Recognition Motif. IDR: Intrinsically Disordered Region. Bottom: schematic depicts the process of meiosis. The period of Rim4 activity is indicated. **(B)** 3D structure of Rim4 predicted by AlphaFold. Yellow indicates low confidence of the prediction; blue indicates high confidence of the prediction. **(C)** GFP-Rim4 forms foci in prophase-I cells under stress conditions. Cells under standard sporulation condition (23°C, 12 h in sporulation medium (SPM), pH 8.5) served as control (Left-top). The tested stress conditions include heat shock (right-top, 37°C for 30 min); acidic solution (left-bottom, pH 5.5) and oxidative condition (right-bottom, 0.5% NaN_3_). White arrows showing the GFP-Rim4 foci formed under stress condition. 30 min heat shock at 37°C followed by 30 min room temperature (23°C) showing the reversible feature of the Rim4 foci. **(D)** Time course (0 min, 5 min, 15 min and 30 min) of GFP-Rim4 foci formation in prophase-I cells at indicated temperatures (25°C, 30°C, 33°C, 37°C, 40°C and 42°C). **(E)** Standard deviation (SD) of the average signal in cells showing in (D). For each condition, 50 cells before (-) and after (temperature as indicated) heat shock were analyzed. See Methods for details. Data were analyzed by Wilcoxon matched-pairs signed rank test. **: p≤0.01; ***: p≤0.001 **(F)** Average foci number per cell over time under each heat shock temperature as showing in (D). All cells (>100) in one field, with or without foci, were analyzed individually for each time points and each condition. **(G)** Representative images of the prophase-I cells harboring mScarlet-Pab1 and GFP-Rim4 with (top) or without (bottom) heat shock (42°C for 10 min). White arrow indicates the foci which mScalet-Pab1 and GFP-Rim4 co-localized. Signals along a white dashed line are analyzed in (H). **(H)** The gray value analysis of the (G). The gray values along the white dashed line in (G) were measured and plotted by the distance. The overlapped peaks showing the co-localized foci of mScarlet-Pab1 and GFP-Rim4. **(I)** FISH of prophase-I cells before (top) or after (middle) 42°C heat shock for 10 min. GFP-Rim4 (green) and mScarlet-Pab1 (red) survived the denaturing process during FISH. mRNAs were detected by Cy5 labeled dT30 (Cy5-dT30) probe (blue). One cell (in yellow box and outlined by white dashed circle) was zoomed in 3 times beside the merge image of heat shocked cells. The gray values along the yellow dashed line in zoomed in image were measured. Lowest value of each channel was considered as background and subtracted from the other data in the same channel, then the highest value was set as 1 and other data were normalized to it. The processed data were plotted by the distance (bottom). Unless otherwise indicated, the scale bar is 5 µm.

Typically, MLOs formed in various cell types or under specific stress conditions, including SGs, share common principles, such as multivalent interactions between proteins and nucleic acids that allow the formation of MLO through liquid-liquid phase separation (LLPS) ^3,4,12–16^. Rim4 comprises three RNA recognition motifs linked by intrinsically disordered regions (IDR) (Fig. 1A; 1B). Surprisingly, the IDR of 289 residues to the C-terminus of Rim4 (IDR_3_), a low complexity domain (LCD), sufficiently drives intracellular Rim4 foci in the nucleus even in the absence of stresses (Fig. 2A). On the other hand, heat-induced foci of Rim4 with C-terminal IDR (IDR_3_) truncated (Rim4[ΔIDR_3_]) are greatly reduced (Fig. 2A), comparing to wild type Rim4 (Fig. 1D). These findings indicate that Rim4 foci can form without mRNAs, primarily driven by IDR_3,_ which plays a role in Rim4 self-assembly ^7^. Consistently, purified recombinant full-length Rim4 conducts phase separation (PS) *in vitro* at its physiological concentration (∼2.5 µM, 150mM NaCl) in the absence of RNAs, favoring low salt concentration, low PH and as low as 2.5% of crowding agent PEG (Fig. 2B; S2A; S2B). In addition, heat (42°C, 10 min) that induced cytosolic Rim4-positive SGs accelerates Rim4 self-assembly *in vitro* (Fig. S2C). These canonical behaviors of proteins conducting PS *in vitro* suggest that Rim4 tends to self-assemble without mRNAs, albeit the physical property and structure of Rim4 self-assembly remain unknown.

**Figure 2.**
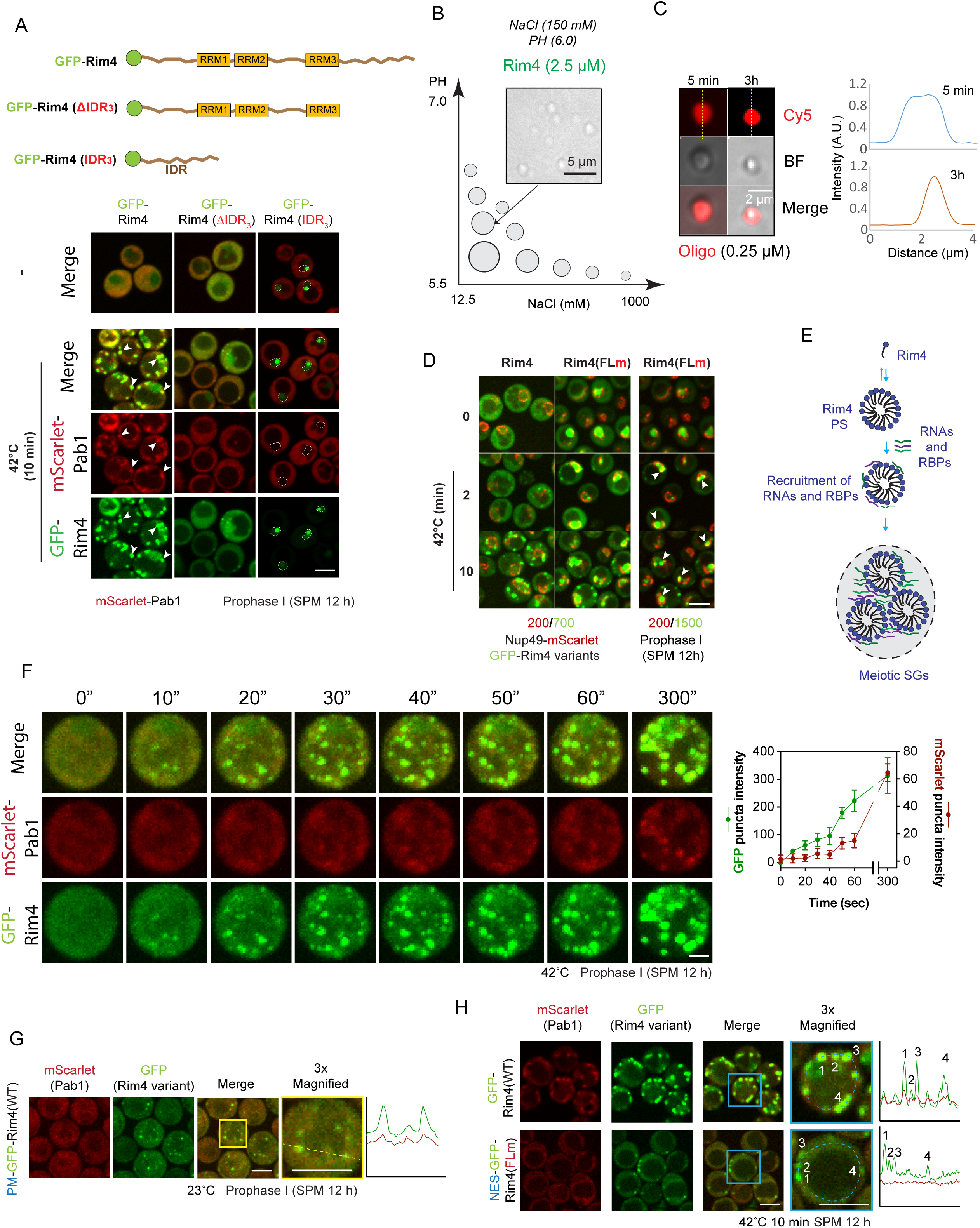
Rim4 self-assembly initiates the formation of meiotic SGs. **(A)** Top: schematic of the labeling and truncation of GFP-Rim4, GFP-Rim4(ΔIDR_3_), and GFP-Rim4(IDR_3_). RRM: RNA Recognition Motif, IDR: Intrinsically Disordered Region. Green round discs represent GFP tag at the N terminus of the proteins. Middle and bottom: Representative images of the GFP-Rim4 (green) variants distribution before (middle) and after (bottom) 42°C heat shock for 10 min. Note that, under heat, the GFP-Rim4 foci formed co-localize with mScarlet-Pab1 (red), while GFP-Rim4(ΔIDR3) reduced foci formation and rare co-localization with mScarlet-Pab1. On the other hand, GFP-Rim4(IDR3) spontaneously forms foci in the nucleus (white dashed circles) without stress, lack of mScarlet-Pab1 co-localization. **(B)** An illustration map showing lower salt concentration and pH value facilitate the phase separation (PS) of recombinant full-length WT Rim4 protein into particles *in vitro* at its physiological concentration (2.5 µM), detected by bright field microcopy. Top center, the representative image of the PS of 2.5 µM Rim4 protein in pH 6.0 SEC buffer (50 mM HEPES-NaOH, pH 6.0; 150 mM NaCl; 2 mM β-mercaptoethanol; 10% glycerol), at 23°C for 5 min. **(C)** 120 nt Cy5 labeled ssDNA oligo (Oligo, red) mimicking the *CLB3* 5’-UTR was recruited to the *in vitro* preformed Rim4 particles. The Rim4 particles was pre-formed as in (B), then mixed with 0.25 µM Oligo. Cy5 and bright field (BF) images were captured after the sample was incubated at 23°C for 5 min and 3 h (left). The Cy5 gray values along the white dashed line in left images were measured and plotted by the distance (right). **(D)** GFP-Rim4(FLm) mutant formed foci in nuclei (white arrow) under heat shock at 42°C. The nuclear envelop is marked by Nup49-mScarlet. The brightness and contrast (B/C) of the left two panels were adjusted in the same way to show the comparable signal level of GFP-Rim4 variants. The B/C of right panel was adjusted to show the foci in nuclei. Please note that in the cells before heat shock (0), GFP-Rim4(FLm) mutant distributed evenly in the nuclei. **(E)** A model of Rim4 initiated SG assembly. Under stress (heat), Rim4 self-assembles into foci with unknown physical property, which recruits RNAs and other RBPs to form SGs. **(F)** left: Still images from a movie of GFP-Rim4 and mScarlet-Pab1 cells under 42°C heat shock, showing that GFP-Rim4 foci forms ahead of mScarlet-Pab1 foci, which eventually co-localize. The time interval was 10 sec. Right: Intensity of both channels on the individual foci were analyzed (n=5). **(G)** Plasma membrane tethered GFP-Rim4 (PM-GFP-Rim4) forms foci without heat shock, co-localize with mScarlet-Pab1. The cell in the yellow box was enlarged 3 times and displayed next to the merge image. The gray values along the yellow dashed line in the enlarged image were measured and plotted by the distance, showing Rim4/Pab1 co-localization. **(H)** NES-GFP-Rim4(FLm) (bottom) co-localize less with mScarlet-Pab1 under 42°C heat shock for 10 min compared to GFP-Rim4 WT (top). The cells in the blue boxes were enlarged 3 times. The gray value along the blue dashed arc were measured and plotted by the distance. The data of 4 foci labeled in the enlarged images were indicated in the plot. Unless otherwise indicated, the scale bar is 5 µm.

As an RBP, the higher order states of Rim4 proteins are relevant to their abilities in binding mRNAs and regulating translation ^7^. *In vitro*, preformed particles of recombinant Rim4 attracts single-stranded DNA (ssDNA) oligos designed to match the 5’UTR element of *CLB3*, a model mRNA substrate of Rim4 ^17^(Fig. 2C). Moreover, the attracted oligos distribute relative evenly in the Rim4 particles within minutes, suggesting the property of these Rim4 particles can efficiently engage mRNAs (Fig. 2C), consistent with that heat-induced cytosolic Rim4 foci recruit mRNAs (Fig. 1I).

In general, cytosolic mRNAs assist SG formation with a structural role ^9^. To examine whether an optimal Rim4/RNA ratio favors Rim4 condensation hypothesizing that RNAs bridge Rim4 molecules, next, we titrated the amount of *CLB3* 5’ UTR oligos from 0.25 µM to 20 µM; surprisingly, the oligos at all concentrations inhibited the formation of Rim4 particles (2.5 µM or 10 µM) (Fig. S2D; S2E). This reminds us that, *in vivo*, mRNA-binding defective Rim4 (IDR_3_) spontaneously forms foci in the nucleus even in the absence of thermal stresses (Fig. 1E).

Notably, although Rim4 almost equally distribute between the nucleus and the cytoplasm ^18^, wild type Rim4 predominantly forms cytosolic foci under heat (Fig. 2D), possibly due to a higher concentration of nascent and available mRNA in the nucleus. To examine this hypothesis, we tested another RNA-binding defective Rim4 mutant, Rim4(FLm). Like Rim4 (IDR_3_), Rim4(FLm) is tethered in the nucleus because that Rim4-mRNA complex assembly in the nucleus is required for efficient Rim4’s export ^19^. Nonetheless, unlike Rim4 (IDR_3_), Rim4(FLm) contains one functional RRM (RRM2) to maintain low level of RNA binding ^19^, preventing its spontaneous self-assembly in the nucleus (Fig. 2D). Strikingly, Rim4(FLm) rapidly form foci in the nucleus, in a sharp contrast to wild type Rim4 (Fig. 2D). Therefore, these results demonstrate that Rim4 self-assembly is an energetically favorable process, counterbalanced by RNA binding.

This finding led us to examine a “seed” model for heat-induced meiotic SGs: Rim4 self-assembly subsequently recruits mRNAs and other SG-resident proteins (Fig. 2E). Pab1, the yeast polyA binding protein, is a major component of yeast SGs. Cytosolic Rim4 foci at initiation stage of meiotic SGs (42°C, 10 sec) exhibited little co-localization with Pab1 (Fig. 2F). In contrast, fully developed Rim4 foci (bigger in size) at later time points of heat shock (42°C, 5 min) recruit Pab1 (Fig. 1G) and Pbp1 (Fig. S1A), another marker of yeast SGs. Because Rim4 interacts with Pab1 in an mRNA-dependent manner ^19^, this finding also suggest that mRNAs are recruited to Rim4-nucleated precursor of SGs at a later time point, consistent with our *in vitro* results showing that preformed self-assembled Rim4 binds nucleic acids (Fig. 2C). Importantly, although the nucleic acids inhibit Rim4 self-assembly (Fig. S2D-E), Rim4 particles upon formation remain stable over time in the presence of nucleic acids (Fig. 2C), thereby allowing the growth of meiotic SGs. Unfortunately, mature meiotic SGs not small Rim4 foci during SG initiation survive a standard FISH procedure, preventing us from examining the timing of mRNA appearance on the Rim4 foci, a topic for future study.

The “seed” model predicts that meiotic SG biogenesis will be reduced in the absence of Rim4 self-assembly (Fig. 2E). In deed, abolishing Rim4 foci formation largely eliminates Pab1-marked cytosolic SGs under 42°C (30 min) (Fig. 2A). On the other hand, Rim4 foci sufficiently recruit Pab1 at 37°C (Fig.S2F), a temperature triggering no Pab1-possitive SGs under vegetative growth conditions ^20^ presumably due to lack of Rim4. Moreover, by anchoring Rim4 to the cytosolic side of plasma membrane, Rim4 (PM-GFP-Rim4) forms foci without stress and recruits Pab1 (Fig. 2G), indicating Rim4 is a primary driver for the biogenesis of meiotic SGs. Notably, PM-GFP-Rim4 carries three intact RRMs for mRNA binding, unlike RNA-binding defective GFP-Rim4(IDR_3_) that forms nuclear foci without recruiting Pab1 (Fig. 2A). Consistent with a role of mRNAs in bridging Rim4 core with other RBPs, the foci of Rim4(FLm) co-localize less with Pab1, comparing with that of Rim4 (Fig. 2H). These results suggest that the thermal meiotic SGs form from C-terminal IDR_3_-mediated Rim4 self-assembly followed by recruitment of mRNAs and other RBPs (e.g., Pab1) via its N-terminal RRMs.

Next, we investigated the regulation of Rim4 self-assembly at the nucleation stage of meiotic SG biogenesis, triggered by thermal stress. The 1,6-Hexonedial (HD) was unable to block heat-induced Rim4 foci formation in meiotic cells (Fig. S3A), suggesting that Rim4 self-assembly is not solely driven by hydrophobicity-based interaction. On the other hand, although Rim4(IDR_3_) primarily mediates Rim4 self-assembly, Rim4(ΔIDR_3_) forms heat-induced foci, albeit with greatly reduced efficiency (Fig. 2A), suggesting that the N-terminal sequence, which contains three RRMs and two IDRs, critically regulates thermal Rim4 foci formation.

About 1/6 of the 713 Rim4 residues are serine (S; 80) or threonine (T; 34), undergoing massive phosphorylation during meiosis with influence on Rim4 self-assembly ^8^ and its interaction with other proteins and RNAs ^18^. Therefore, we speculated that phosphorylation regulates heat-induced Rim4 self-assembly. To identify critical phosphorylation site(s) that regulate heat-induced Rim4 foci, we grouped the S/T residues based on their distribution on Rim4 (R1 to R9; from N- to C-terminus), with all S/T residues in one group simultaneously mutated to alanine (A) or glutamic acid (E). Intriguingly, the E replacement in general reduced Rim4 thermal condensation (Fig. S3B), while R(6-9)-E (Rim4(45E)) almost abolished Rim4 thermal condensation (Fig. 3A), indicating that cumulative phosphorylation of C-terminal IDR_3_ inhibits heat-induced Rim4 foci formation. Strikingly, the R5-A, not R5-E, stimulates Rim4 foci that recruit Pab1 in the absence of heat (Fig. 3B). The R5 region is literally RRM3 of Rim4, which separates the C-terminal IDR_3_ (289aa) from the IDR_2_ (105aa). Conceptually, RRM3 occupied by mRNAs, or Bmh1/2 (through phosphorylated S367/T368) ^18^, would block the potential coordination between the two IDRs in Rim4 self-assembly.

**Figure 3.**
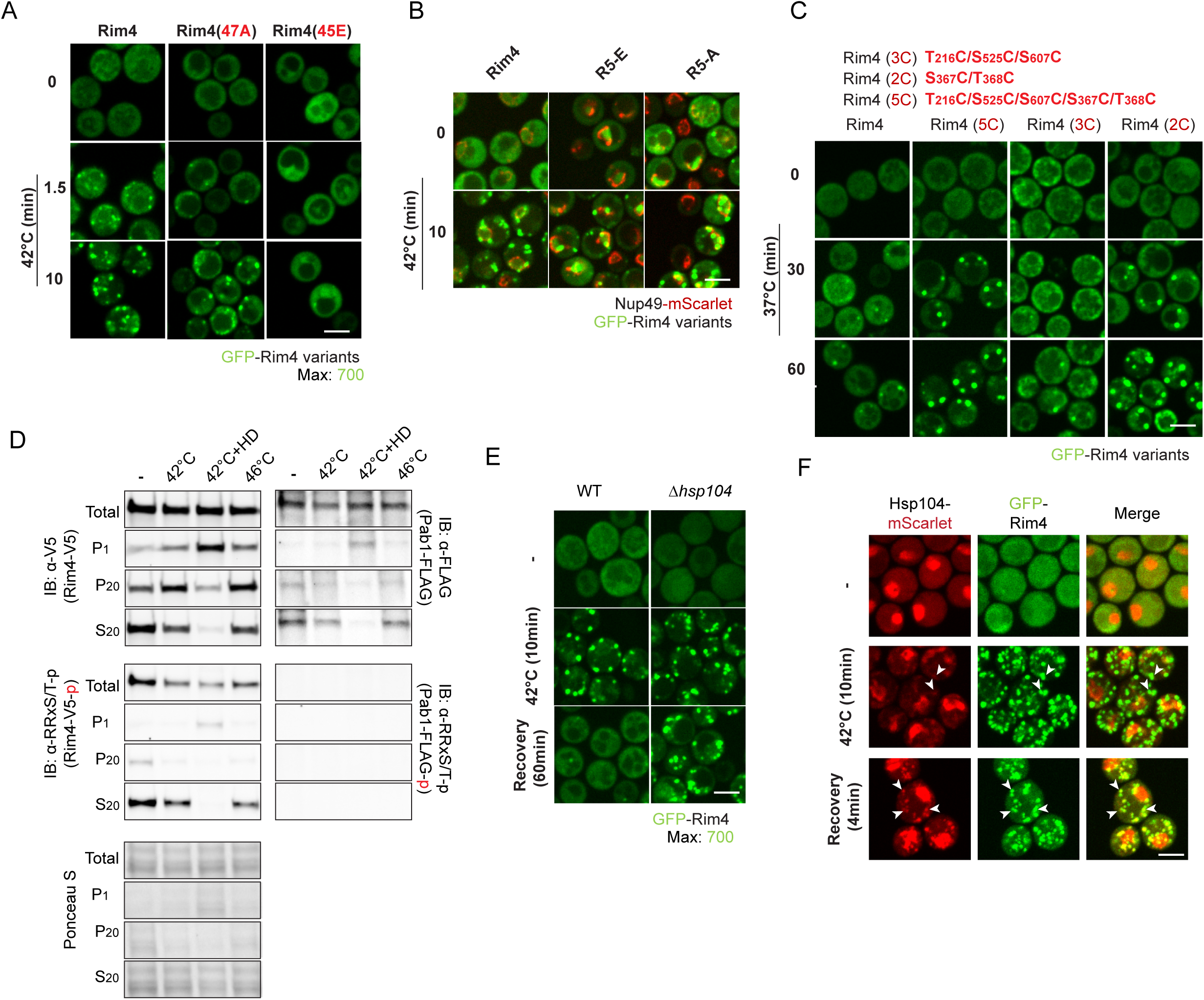
Regulation of the meiotic SGs through Rim4. **(A)** The GFP-Rim4 foci formation in GFP-Rim4 WT, 47A, and 45 E mutation strains under 42°C heat shock at 10 min, showing that GFP-Rim4(45E) mutant almost abolished Rim4 condensation under heat stress. **(B)** GFP-Rim4(R5-A), rather than GFP-Rim4(R5-E) forms foci even without heat (top). Nup49-mScarlet labels the nuclei. Bottom: the GFP-Rim4 variants all form foci under 42°C heat shock for 10 min. **(C)** The representative images of GFP-Rim4, GFP-Rim4(2C) (S367C/T368C), GFP-Rim4(3C) (T216C/S525C/S607C) and GFP-Rim4(5C) (T216C/S367C/T368C/S525C/S607C) behavior before (0) and after 30 min or 60 min heat shock under 37°C. GFP-Rim4(2C) and GFP-Rim4(5C) form foci faster than GFP-Rim4(3C) and GFP-Rim4(WT). **(D)** Immunoblotting (IB) of the whole cell lysates (total) and the fractionation from centrifugations under indicated conditions: with or without 42°C heat shock for 10 min, with 42°C heat shock for 10 min after 5% 1,6-Hexanediol (HD) treatment and with 46°C heat shock for 10 min. IB detects Rim4-V5, PKA phosphorylated Rim4-V5 (Rim4-V5-p) at the RRXS/T sites and Pab1-FLAG. P1: pellet fraction after 1,000 × g centrifugation (10 min); P20: pellet fraction after 20,000 × g centrifugation (10min); S20: supernatant fraction after 20,000 × g centrifugation (10 min). HD: 1,6-Hexanediol. Ponceau S staining serves as even loading control. **(E)** GFP-Rim4 thermal condensates did not disperse in Δhsp104 strain. The cells with WT Hsp104 or Δhsp104 background harboring GFP-Rim4 were heat shocked at 42°C for 10 min, followed by room temperature recovery for 60 min. **(F)** GFP-Rim4 (green) foci formed under heat shock recruit Hsp104-mScarlet (red) during heat shock and recovery (white arrow). The strain harboring GFP-Rim4 and Hsp104-mScarlet were heat shocked at 42°C for 10 min followed by room temperature recovery for 4 min. Unless otherwise indicated, the scale bar is 5 µm.

Besides the Rim4-mRNA complexes, Rim4-Bmh1-Bmh2 is a major intracellular Rim4 complex that contains no RNAs ^18^. Since heat (42°C, 10min) was not sufficient to cancel the inhibitory effect of oligos on Rim4 self-assembly *in vitro* (Fig. S3C), we speculated that the nucleation of cytosolic Rim4 foci derives from dissociation of Bmh1/2 from RRM3, aligning with Bmh1/2’s lack of co-localization with heat-induced Rim4 foci (Fig. S3D). Rim4 harbors four Bmh1/2 binding sites (BBSs) to enable multiple states of Rim4-Bmh1-Bmh2 complex, including the S367/T368 site that resides in RRM3. Next, we found that Rim4(2C), with S367C/T368C to mimic de-phosphorylation state of S367 and T368 to specifically disrupt RRM3-Bmh1/2 interaction, exhibited accelerated thermal SG formation, reminiscent of Rim4(5C), while Rim4(3C) with T216C/S525C/S607C behaved like wild type (Fig. 3C), suggesting that Bmh1/2 suppress Rim4 self-assembly primarily through binding to RRM3. In addition, phosphorylation at Rim4 BBSs was rapidly reduced 10 minutes after moderate heat shock (42°C) (Fig. 3D). Moreover, Rim4 de-phosphorylated at BBSs preferentially resides in the SG-enriched pellet fraction (p20) after centrifugation (20,000 g, 15min) (Fig. 3D). Thus, site-specific de-phosphorylation in RRM3 regulates Rim4 self-assembly through affecting Bmh1/2 interaction. Notably, Rim4(2C) did not form foci like Rim4(R5-A) in the absence of stress (Fig. 3C), suggesting de-phosphorylation at sites other than BBSs in RRM3 might also contribute to Rim4 self-assembly, possibly by interfering RRM3-mRNA interaction. As a conclusion, site-specific phosphorylation (e.g., at S367/T368) in RRM3 and cumulative phosphorylation of IDR_3_ coordinately regulate thermal Rim4 foci formation.

Several studies found that elevated temperatures change certain yeast proteins into reversible SGs ^21^, a dynamic influenced by the severity and length of stresses ^20^. Consistently, heat-induced Rim4 is reversible; the reversibility is affected by the cumulative length of heat exposure (Fig. S4A). Once relieved from short-term thermal stress (42°C, 10 minutes), meiotic SGs marked by Rim4 mainly dispersed in ∼10 minutes, supporting that heat-induced Rim4 foci is distinct from static amyloid aggregation (Fig. S4A). Notably, pab1 disappeared from the SGs before the dispersion of Rim4 during the recovery (Fig. S4A, green arrows), in line with that Rim4 condensation forms a core for the SGs. The recovery kinetics of Rim4(5C) thermal condensates were similar to that of the wild type, indicating that Bmh1/2 did not drive the disassembly (Fig. S4B). Recently, the disaggregation system of Hsp104, Hsp70, and Sis1 (a type II Hsp40), has been shown to mediate rapid dispersion of Heat-induced Pab1 condensates ^20^. To examine the role of this system in Rim4 condensate recovery, we genetically deleted Hsp104 and observed that timely Rim4 dispersion requires heat-shock-induced Hsp104 (Fig. 3E); consistently, Hsp104 was recruited to Rim4 stress condensates (Fig. 3F). Notably, the rapid dispersion of Rim4 thermal condensates (Fig. S4A) implies that Rim4 partial unfolding by the Hsp104 disaggregation system might be sufficient, similar to Pab1 stress granule dispersion ^20^. These findings suggest that the disaggregation system of Hsp104 dissembles Rim4 condensates, conceptually followed by the reformation of Rim4-Bmh1/2 trimeric complex and other forms of Rim4, e.g., Rim4 RNP complex ^18^.

So far, we have identified a meiosis-specific stress granule, i.e., Rim4 stress granule regulated by phosphorylation, the 14-3-3 proteins (Bmh1/2), and the Hsp104 disaggregation system. Intriguingly, due to Rim4, meiotic SGs are formed under temperature as low as 33 °C, as compared to 42°C required for SG formation during yeast vegetative growth. Next, we asked whether the ability of Rim4 self-assembly and thereby the sensitivity of meiotic SGs to heat is critical for meiosis and sporulation. For this purpose, we constructed Rim4(Δ_IDR_) that removes the C-terminal IDR_3_ (Fig. 1D). We also made Rim4-EGFP, aiming to interfere the IDR-driven LLPS by tagging EGFP immediately to the end of IDR (Fig. 4A). As expected, the Rim4(Δ_IDR_) and Rim4-EGFP exhibit impaired self-assembly *in vitro* (Fig. 4B) and foci formation *in vivo* (Fig. 4C, heat shock). Remarkably, Rim4[Δ_IDR_]-EGFP and Rim4-EGFP dramatically reduced meiotic DNA replication (Fig. 4D) and sporulation (Fig. 4E), as compared to Rim4 (EGFP-Rim4). On the other hand, Rim4 variants that form foci spontaneously in cell, e.g., PM-GFP-Rim4 and GFP-Rim4(R5-A) also exhibit severe sporulation defect (Fig. 4F). Notably, PM-GFP-Rim4 and Rim4-GFP are with intact full sequence of Rim4 protein. Thus, the fine-tuned ability of Rim4 self-assembly mediated by IDR is essential for meiosis and sporulation.

**Figure 4.**
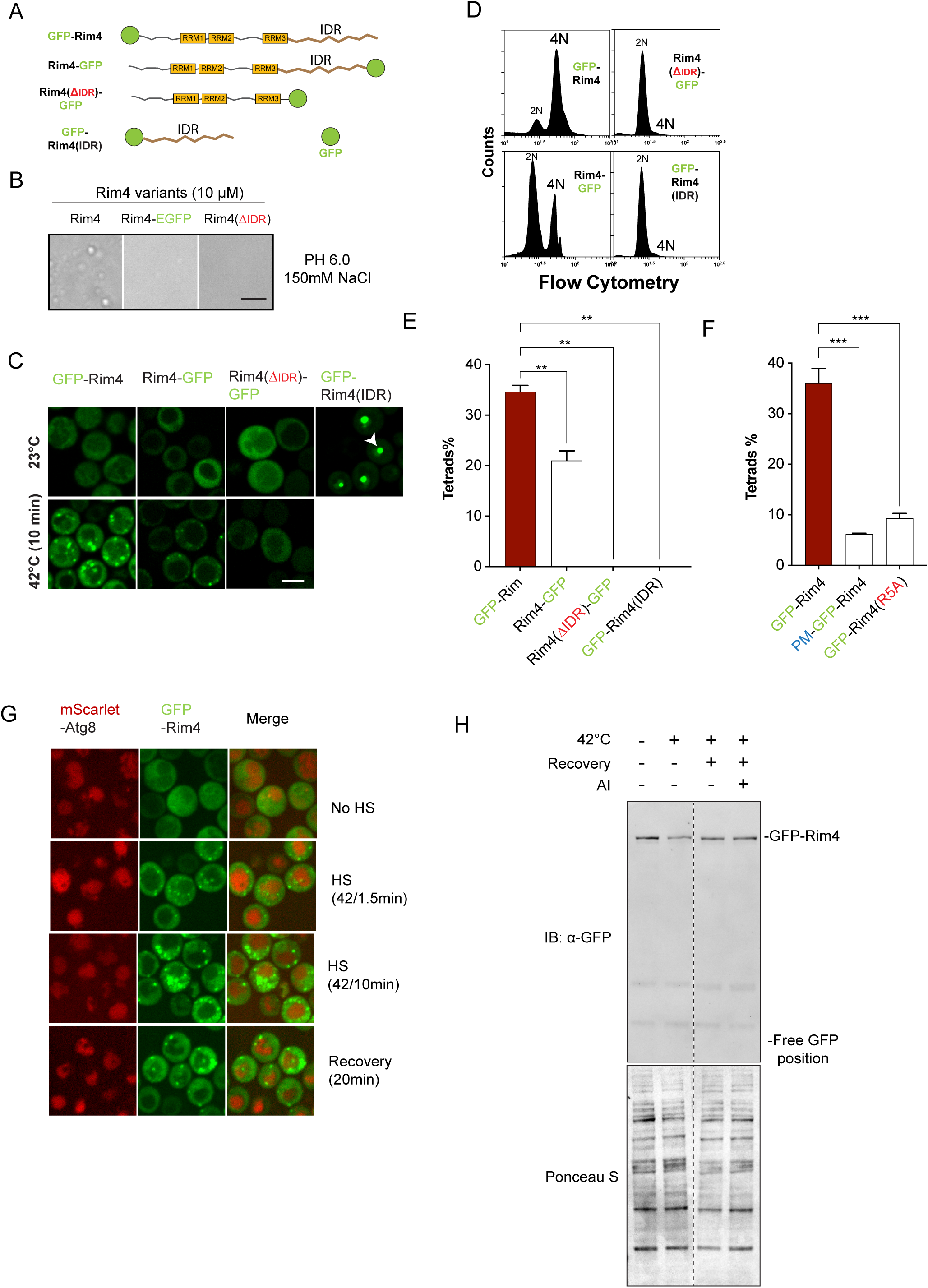
Rim4-positive meiotic SGs resist autophagy. **(A)** Schematic of Rim4 variants with GFP (Green disc) tagged to N terminus or C terminus. RRM: RNA Recognition Motif; IDR: Intrinsically Disordered Region. **(B)** C terminus GFP tagging or IDR3 truncation leads to reduced particle formation (self-assembly) of Rim4 (10 µM, pH 6.0, 5min at 25°C), compared to Rim4 (Rim4-6xHis). **(C)** C terminus GFP tagging or IDR3 truncation of Rim4 inhibits Rim4 foci formation under 42°C heat shock for 10 min. Right, GFP-Rim4(IDR_3_) forms foci in nuclei without heat shock. **(D)** DNA replication of strains harboring GFP-Rim4, Rim4-GFP, Rim4(ΔIDR)-GFP and GFP-Rim4(IDR) at 12 h in SPM. **(E)** Sporulation efficiency of strains harboring GFP-Rim4, Rim4-GFP, Rim4(ΔIDR)-GFP and GFP-Rim4(IDR). **(F)** Sporulation efficiency of strains harboring GFP-Rim4, plasma membrane tethered GFP-Rim4 (PM-GFP-Rim4) and GFP-Rim4(R5A) mutants. Both PM-GFP-Rim4 and GFP-Rim4(R5A) form foci without heat stress. (E) and (F), data from 3 independent experiments were analyzed by unpaired Welch’s t test. **: p≤0.01, ***: p≤0.001. **(G)** Representative images of strain harboring mScarlet-Atg8 and GFP-Rim4 without heat shock (No HS), heat shock at 42°C for 1.5 min, 10 min, and heat shock at 42°C for 10 min followed by room temperature recovery for 20 min. Note no co-localization of GFP-Rim4 foci and mScarlet-Atg8 was observed. **(H)** Immunoblotting of cell lysate (supernatant from 2,000g, 10 min) treated as indicated (42°C heat shock for 10 min, recovery at room temperature for 30 min, AI: 1NM-PP1 to inhibit Atg1-as kinase activity) indicating that GFP-Rim4 is not processed by Autophagy during and after heat shock. Ponceau S staining shows even loading. Unless otherwise indicated, the scale bar is 5 µm.

Next, we investigated how meiosis and sporulation benefit from heat-induced Rim4 self-assembly and meiotic SGs. As a primary suppressor of meiotic translation, Rim4 sequestered a list of mRNAs until its programmed degradation at the end of meiosis I ^17^. We previously reported that autophagy is active during meiosis; therefore, Rim4 (and its target mRNAs) needs to survive autophagy until meiosis II to ensure meiosis and sporulation. So, we examined whether Rim4 condensation, e.g., triggered by thermal stress, is autophagy-resistant. Indeed, autophagy marker Atg8 did not co-localize with heat-induced Rim4 foci in meiotic prophase I cells, respectively, neither during condensation onset nor during recovery (Fig. 4G). Moreover, the level of Rim4 protein in whole cell lysates did not change during short-term heat shock (42 °C, 10 min) and recovery (1 hour); and, importantly, generation of free EGFP by vacuolar digestion of autophagosome-delivered EGFP-Rim4 did not occur (Fig. 4H). Thus, we conclude that Rim4 condensation stabilizes Rim4 under stresses to ensure its programmed degradation at the end of meiosis I, which releases translation for mid-late meiosis and sporulation.

In summary, we reported here that a meiosis-specific RNA binding protein Rim4, which forms amyloid-like aggregates to suppress translation ^7,11^, can nucleate reversible SGs stimulated mainly by heat, presumably stalling meiosis by inhibiting translation to prevent meiotic errors. Although we conducted most intracellular experiments using prophase I cells, the formation of Rim4 SGs occurs whenever Rim4 is present (Fig. S4C) along meiosis. Our findings support a model that meiotic stress granules are regulated by phosphorylation, 14-3-3 protein binding, and the Hsp104 disaggregase system, through Rim4, a core RBP component of the meiotic SGs. Importantly, altering Rim4’s ability to nucleate meiotic SGs impairs meiosis and sporulation even under 25°C, indicating basal level of SG formation might be active and functional during meiosis. Lastly, heat-induced Rim4 foci are distinct from the SDS-resistant amyloid-like aggregates and represent a distinct form of Rim4 self-assembly that recruits mRNAs and other RBPs, such as Pab1 and Pbp1. Future studies should elucidate the composition and function of Rim4-driven meiotic SGs, and how Rim4 converts its protein states to perform functions.

## Method

### Yeast strains and culture

All yeast (*Saccharomyces cerevisiae*) strains in this paper are derivatives of W303 (ade2-1; his3-11,15; leu2-3,112; trp1-1; ura3-1, can1-100).

Deletion strains were created by PCR-mediated knock-out in a parent background with drug resistance (Hyg, Nat, Kan ^22^) or prototrophic markers (Ura3, Leu2, His3, Trp1) as described in previous research ^23^.

Strains with synchronization system (GAL-NDT80/GAL4-ER) were constructed by replacing the NDT80 promoter with the inducible GAL1,10 promoter ^24^.

All strains used in this study carry an analogue sensitive Atg1 mutation (Atg1-as): Atg1(M102G) to allow conditional autophagy inhibition by applying 1NM-PP1, an analogue to ATP ^25,26^.

EGFP-Rim4 variants with RIM4 promoter and terminator were cloned into pRS303 background, and integrated into genomic locus of his3 of the Δrim4 strain. mScarlet-Pab1 and Nup49-mScarlet and Hsp104-mScarlet with their own promoter and terminator were cloned into pRS304 background, and integrated into genomic locus of trp1.

YPD (2% Peptone, 1% yeast extract, 2% glucose), SD complete medium (0.67% Yeast Nitrogen base without amino acids, 2% glucose, complete supplement mixture [CSM]) and YPA (2% Peptone, 1% yeast extract, 2% potassium acetate) were used for vegetative growth of the yeast strains. Standard sporulation medium SPM (0.6% potassium acetate, pH 8.5) was used for yeast sporulation. SD dropout plates (0.67% yeast nitrogen base without amino acids, 2% glucose, auxotrophic amino acids and vitamins, 20% Agar) were used for yeast transformation/dissection selection.

### Sporulation

The sporulation process was described in previous publications ^11,18,27^. A single colony of a yeast strain was picked up and spread on a YPD plate. The cells were growing under 30°C for o/n (about 24 h) until cells formed a lawn. Collect the cells by scratching the plate and suspend them in YPA medium (OD_600_ = 0.3) and grown for 14 h at 30°C. Cells were then spun down, washed with SPM twice and resuspend in SPM to final OD_600_ = 2.

For the strains with synchronization system (GAL-NDT80/GAL4-ER) were released from the prophase-I arresting by addition of 1 µM β-estradiol (Sigma E2758-1G) at 12 h in SPM.

At 60 h in SPM, count the total cell number and the number of cells containing four spores (tetrads), respectively, with a hemocytometer. Calculate the tetrads percentage by divide the tetrads number by the total cell number. More than 300 cells were counted for each strain. Data from at least 3 independent experiments were processed by GraphPad Prism 10, plotted as Mean ± Standard error (SE) and analyzed by unpaired Welch’s t test.

### Living cell image

200 µL cells were collected by spun down. Resuspend with 20 µL original media. Drop 5 µL to a wide No. 1½ glass slides (Corning, 2975-246) or Smart Substrate of VAHEAT (Interherence GmbH) (see below). The cells were covered by a piece (∼ 5 mm × 5 mm × 5 mm) agar containing the same media, and immediately observe with fluorescent microscopy (FM). The microscope is a Zeiss Axio Observer with Spinning disk Confocal (Yokogawa Spinning Disk Confocal CSU-W1) supplied with Hamamatsu Orca-Fusion sCMOS camera and a Zeiss Plan Apochromat 63× /0.9-NA oil-immersion objective. Exposure time was 250 ms for both mScarlet (laser: 561 nm, power: 10%, filter: 617 / 71 nm,) and GFP (laser: 473 nm, power: 20%, filter: 525 / 50 nm) channels. Images were captured with SlideBook 2022 software (Intelligent Imaging Innovations). Images of 5 planes z stacks, 2.5 µm per plane, were taken for each channel. For static images, 3 fields (188.94 μm × 191.64 µm) containing more than 300 cells were taken for each sample. For movies, 1 field containing more than 100 cells were taken for each sample, each time point.

The original images were exported to Tagged Image File Format (TIFF) for each channel, each time point. For static images, the images were split for the best focus, pseudo-colored and analyzed by ImageJ; for movies, the images in a series were z projected, pseudo-colored and concatenated into TIFF for analysis or Audio Video Interactive (AVI) format for movie output.

### FM with Heat shock

Two ways were used to conduct heat shock to the cells for FM experiments:

### Movies of cells under heat shock with VAHEAT

200 µL cell were concentrated to 20 µL as described above. Insert the Smart Substrate into the VAHEAT adaptor, connect the connector head and place the unit in the microscope stage. Drop 5 µL cell to the center of Smart Substrate. Cover the cell drop with a piece (∼ 5 mm × 5 mm × 5 mm) agar containing the same media. Set the profile of heat shock in the VAHEAT UI software (Interherence GmbH). A typical heat shock profile including: 10 sec count down at 25°C (RT), 10 sec, 30 sec or 2 min at RT for the first two images interval, 0.1 sec (as short as possible) heating up to specific temperature, specific time for heat shock at the specific temperature, 0.1 sec (as short as possible) cooling down to 25°C, and optionally, recover at 25°C.

For the experiments studying the early time points of the heat shock, set the time interval of the movie at 10 sec. Also set the second step of the heat shock profile at 10 sec. After the cells were covered by agar, find a field which containing at least 100 cells. Start the heat shock profile. After the 10 sec count down, start take the movie. So that the first image is captured under 25°C, and the second image is captured when heat shock started.

For the experiments studying the full process of heat shock, set the time interval of the movie at 30 sec. Also set the second step of the heat shock profile at 30 sec. After the cells were covered by agar, find a field which containing at least 100 cells. Start the heat shock profile. After the 10 sec count down, start take the movie. So that the first image is captured under 25°C, and the second image is captured when heat shock started.

For the experiments studying both the heat shock and recovery, set the time interval of the movie at 2 min. Also set the second step of the heat shock profile at 2 min. After the cells were covered by agar, find a field which containing at least 100 cells. Start the heat shock profile. After the 10 sec count down, start take the movie. So that the first image is captured under 25°C, and the second image is captured when heat shock started.

For the experiments studying the recovery, set the time interval of the movie at 30 sec. Also set the second step of the heat shock profile at 30 sec. After the cells were covered by agar, find a field which containing at least 100 cells. Start the heat shock profile. After the heat shock, start to capture the movie.

### Static image of cells under heat shock with PCR machine

200 µL cells were concentrated to 20 µL as described above. Transfer the cells into a PCR tube sit in the PCR machine. Set the PCR machine to specific temperature in the “incubation” mode, and the lid temperature 5 degree higher to prevent evaporation. After the heat shock, transfer the PCR tube in an insulation bottle containing the same heat shock temperature water to the microscope, with the VAHEAT on and set to the same heat shock temperature. Then, drop 5 µL of the cells to the pre-heated Smart Substrate, and capture images as described above.

### Signal distribution analysis of the FM movies

Outline individual cells at the first time point (before heat shock) with ImageJ’s Freehand Selections tool, measure the mean and standard deviation (SD) of the intensity of this channel. Move to the time point after 10 min heat shock, and measure the same parameters for the same area of the same cell. 50 cells were measured for each sample. Data were processed with GraphPad Prism 10, plotted as scatterplot, displaying mean ± SE, and analyzed by Wilcoxon matched-pairs signed rank test.

### Puncta counting of the FM movies

Extract the frames showing 0 min, 5 min, 10 min, 15 min, 20 min, 25 min and 20 min heat shock of the movie in ImageJ. Count the puncta in each cell. If there are no puncta in one cell, then count as zero. For each image, count all the cells in the field (120∼200 cells). Data were processed with GraphPad Prism 10, plotted as violin plot showing all points and calculate the mean and SD. Plot the mean and SD of groups in comparison (e.g., different temperature) as broken line chart with GraphPad Prism 10, showing the average puncta number per cell at different temperature, different time points.

### Gray Value analysis of individual cells or condensation

Pick a representative cell or condensation image. Draw a line (or curve to avoid vacuole) through the interested structure (e.g., the foci in cell, or *in vitro* preformed condensation). The gray values along the line or curve were measured in ImageJ, and the data were plotted by the distance with GraphPad Prism 10.

### Protein recruitment analysis

Extract the frames showing 0 sec, 10 sec, 20 sec, 30 sec, 40 sec, 50 sec, 60 sec and 300 sec heat shock of the movie in ImageJ. Pick a representative cell for analysis. 5 foci were selected by a 3 pixels (0.51 µm) diameter circle in each frame, referred by GFP-Rim4 (green) channel. the signals of the foci were measured in both channels. For the start time point, 5 random area were selected to measure. Additionally, signals of 5 background areas were also measured as described above. The average background signal was subtracted from the foci signals in the same frame. The results were plotted by the heat shock time with GraphPad Prism 10.

### Immunoblotting

1.5 OD_600_ cells were spun down. Resuspend in 20 µL SPM containing 10 µM PMSF. Sit at room temperature for 5 min. Spin down and store the pellets in -80°C until the gel running.

Resuspend the pellets in 100 µL 2× SDS loading buffer (125 mM Tris-HCl, pH 6.8; 4% SDS; 0.1% BPB; 20% Glycerol; 10% β-mercaptoethanol; 2× cOmplete Protease Inhibitor Cocktail [Roche, 11873580001]; 2 ng/µL Pepstatin; 2 µM PMSF) and heat at 70°C for 10 min. Spin down at maximum speed (14,000 rpm) for 5 min. Load 10 µL supernatant to 4∼20% gradient PAGE gels (26-well: Criterion® Tris-Glycine[TGX] Stain-Free Gels [Bio-Rad, 5678095]; 15-well: SuperPAGE™ Bis-Tris Gels [GenScript, M00657]). Run at 200 V for 30 min. Subsequentially, the samples were electroblotted onto nitrocellulose (NC) membranes (Bio-Rad, 1620115) with the Trans-Blot semi-dry transferring cell (Bio-Rad, 1703940).

The NC membrane blotted with the protein samples were stained in 0.1% Ponceau S solution (0.1% Ponceau S, 5% acetic acid) at RT for 3 min for the evidence of even loading. Then the membrane was blocked in 1% BSA in TBST (20 mM Tris-HCl, pH 7.5; 150 mM NaCl; 0.5% Tween-20) at RT for 45 min. Next, incubate the membrane in primary antibody diluted in TBST containing 1% BSA (Mouse-anti-V5 antibody[Fisher, PIMA515253]: 1:2,500; Mouse-anti-FLAG antibody [Sigma-Aldrich, F3165]: 1:5,000; Mouse-anti-GFP antibody [Roche, 11814460001]: 1:5,000; Rabbit anti-Phospho-PKA substrate [RRxS*/T*] antibody [Cell Signaling Technologies, 9624S]: 1:2000; Rabbit-anti-Pgk1: 1:10,000) at 4°C for o/n. After wash with TBST for 3 times at RT, the membrane was incubated in secondary antibody diluted in TBST containing 1% BSA (StarBright B700 conjugated Goat-anti-Mouse antibody [Bio-Rad, 12004158]: 1:10,000; Fluor Alexa 488 conjugated Goat-anti-Rabbit antibody [Thermo-Fisher, A11034]: 1:10,000) at RT for 45 min. Finally, wash with TBST for 3 times and capture the IB images with ChemiDoc MP Imaging System (Bio-Rad, 12003154).

### Recombinant protein purification

Rim4 and mCherry-Pab1 open reading frames (ORFs) were cloned into pET29b(+) background, with the 6× His tag at the C terminus; GFP-Rim4 ORF were cloned into pET28a(+) background, with the 6× His tag at the N terminus. BL21(DE3) *E. coli* strains harboring the pET plasmids were induced by IPTG as described previously^28^ For Rim4 and GFP-Rim4, continue shake at 16°C for overnight (∼14 h); for mCherry-Pab1, shake at 37°C for additional 4 h.

For Rim4 and mCherry-Pab1, collect the bacteria and lyze with a High-Pressure Cell Press Homogenizer (Avestin Emulsiflex-C5) in lysis buffer (BLB: 50 mM Tris-HCl, pH 8.0; 300 mM NaCl; 10 mM MgCl_2_; 10 mM Imidazole; 10% Glycerol; 5 mM β-mercaptoethanol; 1 mM PMSF; 1× cOmplete Protease Inhibitor Cocktail [Roche, 11873580001]; 1 ng/µL Pepstatin). The lysate was treated with 0.1 mg/mL DNase I and 0.1 mg/mL RNase A and 0.1% Triton X-100 at 4°C for 30 min on a rotator. Clearing the lysate by spinning down at 30,000 × g for 1 h at 4°C. Apply the supernatant to 1 mL bed volume of NTA-Ni column (QIAGEN, 30230), and wash with BLB supplied with gradually increased concentration of Imidazole (10 mM, 25 mM and 50 mM) and gradually reduced concentration of NaCl (500 mM, 300 mM and 150 mM). The protein was then eluted with elution buffer (50 mM Tris-HCl, pH 8.0; 150 mM NaCl; 250 mM Imidazole; 10% glycerol and 5 mM β-mercaptoethanol). The eluant was subsequentially separated by Superdex 200 Increase 10/300 GL column (GE healthcare) equilibrated in Size Exclusive Buffer (SEC Buffer: 50 mM Tris-HCl, pH 8.0; 150 mM NaCl; 10% Glycerol; 2 mM β-mercaptoethanol).

Fractions were analyzed by SDS-PAGE followed by Coomassie Bright Blue R-250 (CBB) staining. The purist peak fractions were pooled and concentrated with a Amicon Ultra centrifugal filter (Sigma). The purified proteins were snap frozen in liquid nitrogen. The concentration of the protein was determined by Bradford Assay^29^ and the purity of the protein were qualified by SDS-PAGE followed by CBB staining.

For GFP-Rim4, use high salt buffers through the process, due to the protein’s propensity of condensation/aggregation. i.e., substitute all the NaCl with 1 M NaCl. The other process is as same as described above.

### *In vitro* condensation and microscopy

The Rim4 protein were diluted in SEC buffer with indicated pH, salt concentration, and/or additives (e.g., PEG, ssDNA, 1,6-Hexanediol, RNase A, Pab1 protein, etc.) to the specified concentration (2.5 µM or 10 µM). The diluted protein samples were then incubated at different temperature, before load to glass slides and capture images under microscope. No cover glass or agar was applied to avoid any possible destruction of the condensates. A PCR tube lid was used to cover the drop to prevent evaporation during long term (3 h) incubation on slide.

### DNA replication analysis with flowcytometry

Unless otherwise specified, the cells arrested at Prophase-I (12 hr in SPM) were collected by centrifugation and fixed in 70% ethanol for overnight. Spin down and resuspend in 50 mM Sodium Citrate. Ultrasonicate the cells at 30% power for 15 sec. Spin down and resuspend in 50 mM Sodium Citrate supplied with 0.5 mg/mL RNase A. Incubate at 37°C for overnight. Spin down and resuspend in 50 mM Sodium Citrate supplied with 2.5 µM SYTOX Green. Perform flowcytometry with BD FACSCalibur flow cytometer. Data were analyzed by Flowing Software 2.5.1 (University of Turku, Finland). The living cells were gated based on front scattering (FSC) and side scattering (SSC). Generate a histogram depicting cell counts in the Green (488-530/30 nm) channel.

### Fluorescence *in situ* hybridization (FISH)

FISH assay was conducted as described previously ^30^. Briefly, collect 3.5 OD_600_ cells by centrifugation. Immediately resuspend in 2 mL fixation buffer (∼3% formaldehyde in SPM). Rotate at RT for 20 min followed by in 4°C for overnight. Wash with ice-cold 1× Buffer B (1.2 M D-sorbitol in 100 mM KPi [Potassium Phosphate buffer, pH 7.5]) and then store the pellets in -80°C until the next step.

Digest the cell wall in 430 µL 1× Buffer B containing 0.116 mg/mL Zymolyase at 30°C until ∼80% of cell’s cell wall was digested (∼15 min). Spin down at 380 × g for 5 min. Remove the liquid completely by pipetting. Gently wash twice with 1 mL 1× Buffer B. Resuspend in 1 mL 70% Ethanol, and incubate in RT for 4 h. Transfer the cells to low-binding microtubes (USA Scientific, 1415-2600). Collect the cells by spinning down at 380 × g for 5 min. Treat the pellets with 1 mL formamide wash buffer (FWB: 10% formamide, 2× SSC[300 mM NaCl, 30 mM Sodium Citrate]) at RT for 20 min. Then, in darkness, the cells were incubated in the hybridization solution (0.2 µM Cy5 labeled probe in hybridization buffer [0.1 g/mL dextran sulfate sodium; 10% deionized formamide; 2× SSC]) at 30°C on rotator for overnight.

Pellet the cells at 380 × g for 3 min. Gently incubate the cells with 1 mL FWB at RT for 30 min. Pellet again and remove the liquid as much as possible. Resuspend in 20 µL DEPC-PBS and proceed with FM.

The GFP and mScarlet protein tags can partially survive denaturation in the FISH assay, we can detect the protein signal and the mRNA signal in the same experiment. Exposure time was 250 ms for GFP channel (Laser 473 nm, power: 30%, filter: 525/50 nm); 500 ms for mScarlet channel (Laser: 561 nm, power: 10%, filter: 617/73 nm); 250 ms for Cy5 channel (Laser: 640 nm, power: 10%, filter: 692/40 nm). Z-stack of 11 planes, 1 µm per plane. Capture 3 fields (188.94 µm × 191.64 µm) containing at least 300 cells for each sample.

To detect the total mRNAs in the cells, use Cy5 labeled dT_30_ (Cy5-TTTTTTTTTTTTTTTTTTTTTTTTTTTTTT) as probe, which can specifically bind to the Poly(A) tails of the mRNAs.

## Acknowledgement

We thank Vladimir Denic (Harvard Unisersity, Boston, MA) for reagents and protocols. We thank J. Seemann, J. Friedman, M. Henne, and members of the Wang lab for comments on the project.

This work was funded by grants from the National Institutes of Health (R01GM133899) and from the Welch Foundation (I-2019-20190330), and funding from Nancy Cain and Jeffery A. Marcus Scholar in Medical Research, in honor of Dr. Bill S. Vowell, to F.W..

## Author Contribution

R.Z., S.L., W.F. and F.W. conceptualized and designed the project. R.Z., S.L., W.F., S.Q., A.J.C. and F.W. performed experiments. R.Z. and S.L. quantified and analyzed the data.

R.Z. and F.W. wrote the manuscript and interpreted the data with the help of critical reviews from all coauthors

---

**Figure S1. Characterization of heat-induced Rim4 foci.**

(**A**) Representative FM images showing GFP-Rim4 (green) and mScarlet-Pbp1 (red) co-localized under heat shock at 42°C for 10 min in prophase-I cells. **(B)** Representative FM images showing GFP-Rim4, GFP-Rim4 with single (ΔpolyN1 and ΔpolyN2) or double (ΔpolyN12) poly asparagine (polyN) region deletion forms foci under 42°C heat shock for 10 min. **(C)** GFP-Rim4 thermal condensates formed under 42°C heat shock for 10 min partially disassembled by 0.5% Triton X-100 treatment at room temperature for 10 min, and fully disappear after 2% SDS treatment at room temperature for 10 min. Unless otherwise indicated, the scale bar is 5 µm.

**Figure S2. Characterization of Rim4 self-assembly *in vitro* and in cells.**

**(A)** Rim4 recombinant protein forms condensates at low salt, low pH condition. Rim4 recombinant protein was diluted to 2.5 µM in SEC buffer (50 mM HEPES-NaOH at indicated pH; NaCl at indicated concentration; 2 mM β-mercaptoethanol; 10% glycerol) and incubated at room temperature for 5 min or 80 min (as indicated, for pH 5.5, 1000 mM NaCl only) before bright field microscopy. **(B)** PEG stimulates phase separation of Rim4 recombinant protein into particles. 10 µM Rim4 were incubated with crowding reagent PEG3350 at indicated concentration in pH 6.8 SEC containing 150 mM NaCl. **(C)** Heat (42°C) stimulates phase separation of Rim4 recombinant protein into particles. 2.5 µM Rim4 were incubated in pH 6.0 SEC containing 300 mM or 500 mM NaCl at indicated temperature for 10 min. **(D-E)** ssDNA inhibits Rim4 recombinant protein forming condensates. 120 nt ssDNA oligo mimicking *CLB3* 5’UTR at indicated concentration were co-incubated with 2.5 µM (D) or 10 µM (E) Rim4 protein in pH 6.0 SEC containing 150 mM NaCl at 22°C for 5 min or overnight (o/n) as indicated in (D). **(F)** GFP-Rim4 (green) foci recruits mScarlet-Pab1 (red) under 37°C heat shock condition. The cells were heat shocked at 37°C for 30 min before FM microscopy. White arrows indicate the co-localization of GFP-Rim4 and mScarlet-Pab1. Unless otherwise indicated, the scale bar is 5 µm.

**Figure S3. Phosphorylation and Bmh1/Bmh2 regulates heat-induced Rim4 foci formation**

**(A)** 1,6-Hexandiol (HD) did not dissolve the Rim4 foci stimulated by heat shock. The cells with or without 5% HD treatment at RT for 10 min were heat shocked at 42°C for 10 min before FM microscopy. **(B)** Representative FM images of the indicated GFP-Rim4 (green) variants forming foci after 42°C heat shock for 1.5 min or 10 min. Nup49-mScarlet (red) serves as nuclear membrane marker. **(C)** The inhibition of in vitro Rim4 self-assembly by ssDNA cannot be cancelled by increasing temperature to 42°C. ssDNA mimicking *CLB3* 5’UTR at indicated concentration were co-incubated with 2.5 µM Rim4 recombinant protein in pH 6.0 SEC containing 150 mM NaCl at indicated temperature for indicated time before bright field microscopy. **(D)** Representative FM images showing that GFP-Rim4 thermal condensation lacks efficient interaction with the 14-3-3 proteins Bmh1 and Bmh2. Cells were heat shocked at 42°C for 10 min before FM microscopy. Unless otherwise indicated, the scale bar is 5 µm.

**Figure S4. The assembly and disassembly of meiotic SGs center on Rim4.**

**(A)** The length (times) of heat exposure affects meiotic SGs disassemble during recovery. The cells were heat shocked and placed under recovery at room temperature as indicated. White arrows indicated the foci of GFP-Rim4 and mScarlet-Pab1 colocalization; green arrows indicated the foci of GFP-Rim4 only, without mScarlet-Pab1. Notably, Pab1 leaves SGs before Rim4 foci fully disperse. **(B)** GFP-Rim4(5C) foci disassembly follows similar pattern as that of GFP-Rim4WT. Both cells were heated shocked at 42°C for 10 min and recovered for 30 min and 60 min before FM microscopy. **(C)** The formation of meiotic SGs (in SPM at indicated time) showing no meiotic stage restriction. Cells in SPM for indicated time were collected for FM microscopy without (top) and heat shocked at 42°C for 30 min (bottom).

