## Supplementary figures and images for "Rim4 is a Thermal Sensor and Driver of Meiosis-specific Stress Granules"

### Supplementary Figure S1

Zhang et al. Figure. S1

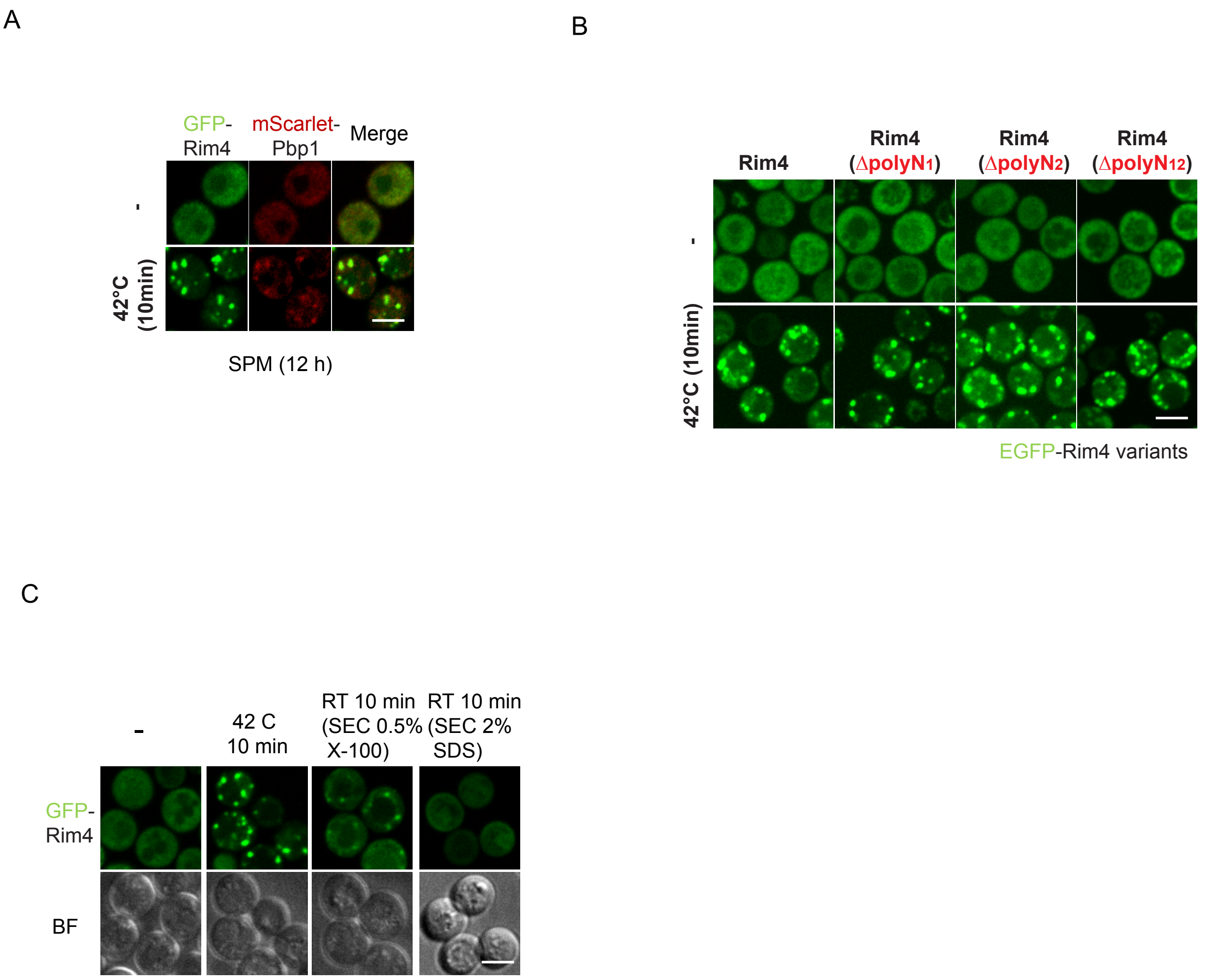

### Supplementary Figure S2

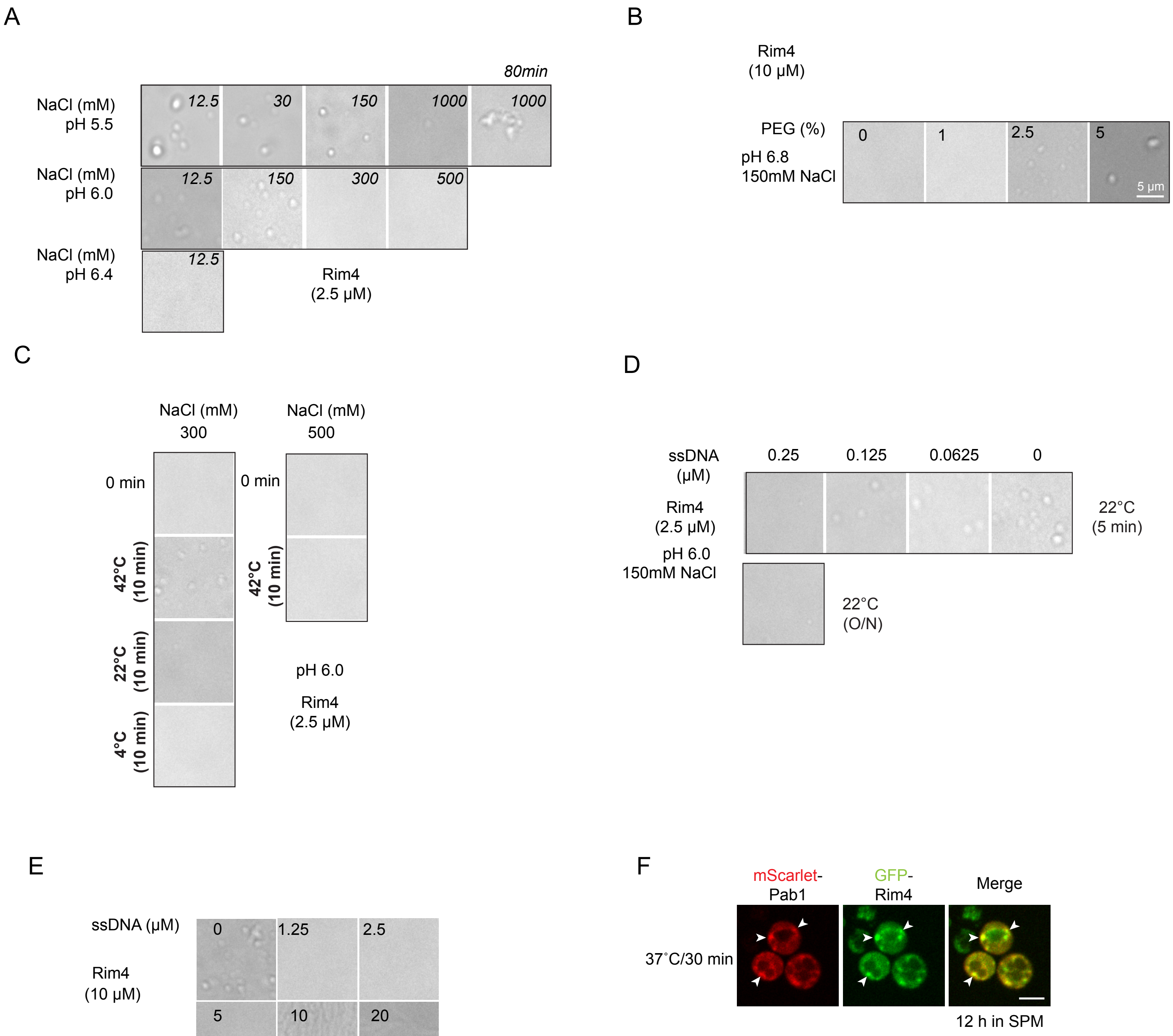

### Supplementary Figure S3

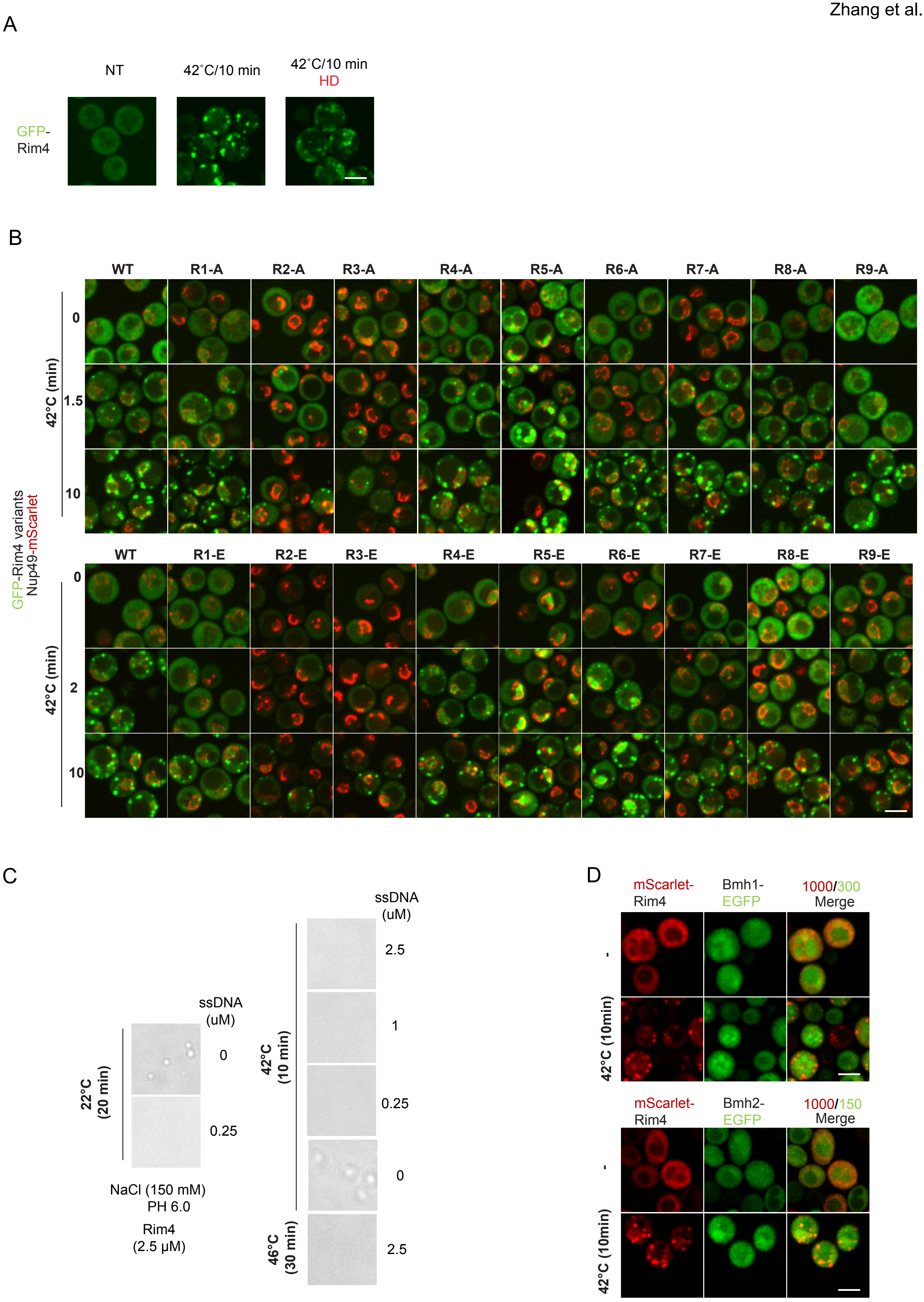

### Supplementary Figure S4

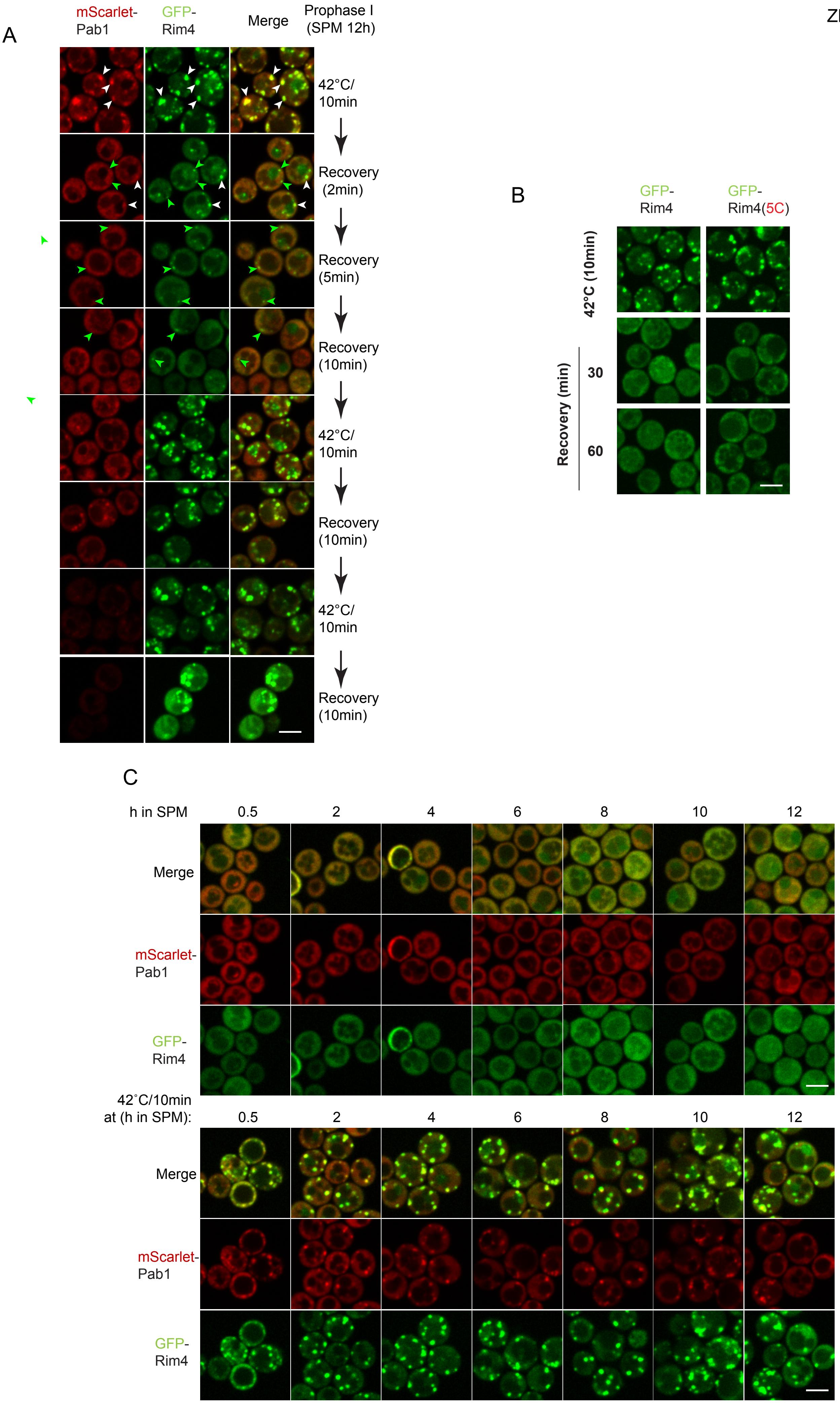
